# Longevity interventions temporally scale healthspan in *Caenorhabditis elegans*

**DOI:** 10.1101/2021.05.31.446397

**Authors:** Cyril Statzer, Peter Reichert, Jürg Dual, Collin Y. Ewald

## Abstract

Human centenarians and longevity mutants of model organisms show lower incidence rates of late-life morbidities than the average population. However, whether longevity is caused by a compression of the portion of life spent in a state of morbidity, *i*.*e*., “sickspan,” is highly debated even in isogenic *C. elegans*. Here, we developed a microfluidic device that employs acoustophoretic force fields to quantify the maximum muscle strength and dynamic power in aging *C. elegans*. Together with different biomarkers for healthspan, we found a stochastic onset of morbidity, starting with a decline in dynamic muscle power and structural integrity, culminating in frailty. Surprisingly, we did not observe a compression of sickspan in longevity mutants but instead observed a temporal scaling of healthspan. Given the conservation of these longevity interventions, this raises the question of whether the healthspan of mammalian longevity interventions is also temporally scaled.

## Introduction

The continuously growing elderly population is projected to result in 1.5 billion people above the age of 65 globally by 2050 (Nations, 2019). This poses a significant challenge since old age is the major risk factor for developing cancer, dementia, cardiovascular, and metabolic diseases (Partridge et al., 2018), especially since people suffer for approximately 20% of their lifespan from one or multiple of these chronic illnesses, which are themselves accompanied by other late-life disabilities (Partridge et al., 2018). Current estimates indicate that delaying the onset of these chronic diseases by one year would save $38 trillion in the US alone (Scott et al., 2021). Therefore, major research efforts are dedicated to understanding how to increase the time spent in good health (*i*.*e*., healthspan) and to postpone and compress the time spent suffering from age-related pathologies and chronic diseases (*i*.*e*., sickspan) (Kaeberlein, 2017; Kennedy et al., 2014; Olshansky, 2018; Partridge et al., 2018).

People that are more than one hundred years old, so-called centenarians, display a delayed onset and a lower incidence rate of late-life morbidities compared to people in the age bracket of 80 to 89 years (Ailshire et al., 2015; Andersen et al., 2012; Evans et al., 2014; Evert et al., 2003; Ismail et al., 2016; Kheirbek et al., 2017). Genome-wide association studies have shown associations between the exceptional longevity of centenarians and aging-related genes identified in model organisms (Kenyon, 2010; López-Otín et al., 2013; Partridge et al., 2018). Mutations in genes that promote longevity in model organisms, such as *C. elegans*, have been instrumental in identifying mechanisms that promote healthy aging (Kenyon, 2010; López-Otín et al., 2013; Magalhães et al., 2017; Partridge et al., 2018).

A recent study has questioned this approach of using *C. elegans* longevity mutants to gain insights for promoting healthy aging or mechanisms that prolong healthspan (Bansal et al., 2015). Using four matrices (resilience to heat and oxidative stress, voluntary movement, and swimming performance) to assess the “health” status of aging *C. elegans*, they found that four commonly used longevity mutants outperformed wild type at any given timepoint at older ages, consistent with previous reports. However, compared to their maximum lifespan, longevity mutants displayed an increased sickspan-to-healthspan ratio compared to wild type (Bansal et al., 2015). Other studies have not observed an increase of sickspan in long-lived *C. elegans* mutants, except in the case of lower mobility or movement scores for the insulin/IGF-1 receptor longevity *daf-2(e1370)* mutants (Hahm et al., 2015; Huang et al., 2004; Podshivalova and Kerr, 2017; Stamper et al., 2018). Part of the “prolonged sickspan” based on the motility of these *daf-2(e1370)* mutants was attributed to lack of behavioral exploration linked to *odr-10* gene expression (Hahm et al., 2015) and improper dauer-like quiescence behavior (Ewald et al., 2015, 2018; Gems et al., 1998; Hess et al., 2019; Podshivalova and Kerr, 2017). Although all these studies showed that sickspan is not increased in longevity mutants, the question remained about how healthspan changes when the lifespan is extended. We hypothesized that using other health matrices independent of voluntary or behavioral influences, such as physical properties of muscular strength, which is one of the best predictors for all-cause mortality in humans (Leong et al., 2015), we might be able to quantify the health trajectory of *C. elegans* longevity mutants.

Here we confirm that voluntary movement during aging declines, and this fragility is not extended in longevity mutants, except mildly in *daf-2* mutants, using high-resolution lifespan and movement measurements on plates. We developed a novel microfluidic device and applied acoustophoretic force fields to quantify the maximum force and power of *C. elegans*. Using a high-frequency and high-power acoustic force field, it becomes possible to set up a contactless, constant in time, and uniform force field acting along the whole *C. elegans* body. Therefore, this force field challenges swimming *C. elegans* in a similar way body-weight exercises do for humans in a gravity field. Furthermore, applying the acoustic field stimulated a swimming response of resting *C. elegans*. All longevity mutants showed delayed onset of the decline in maximum force and dynamic power during aging. We observed heterogeneity between individuals across all genotypes in the onset of age-related phenotypes, several correlated phenotypes, and a time-dependent occurrence of multiple disabilities. However, we did not find a compression of sickspan, but rather a temporal scaling of healthspan relative to their maximal lifespan across genotypes.

## Results

### Voluntary movement healthspan is proportionally increased by longevity interventions

To obtain highly quantitative data on lifespan and healthspan, we used a lifespan machine (Stroustrup et al., 2013). Here, we defined the “voluntary movement healthspan” as the time spent fast crawling and the “voluntary movement sickspan” as the time spent slow crawling or displaying minimal posture changes (see Materials and Methods for detailed definition). We chose *eat-2(ad1116)* as a genetic model for dietary restriction-like mediated longevity, *glp-1(e2141)* as a genetic model for germ-stem-cell-less-mediated longevity, *daf-2(e1368)* and *daf-2(e1370)* as genetic models for reduced insulin/IGF-1 signaling mediated longevity. We cultured all animals at the same temperature (15°C) and in the same environment with the same food source, except *glp-1* that underwent a brief temperature upshift during development as in preparation for the lifespan assay. To avoid dauer-specific traits that occur in reduced insulin/IGF-1 signaling mutants (Ewald et al., 2018) and to avoid pathogenicity from a bacterial food source (Podshivalova and Kerr, 2017), lifespans were run at 15°C on heat-killed bacteria. Thus, the experimental setup was designed to offer optimal conditions and was kept identical while *C. elegans* genotypes were varied.

As expected, we measured a significant increase in lifespan for these long-lived mutants compared to wild type (Figure 1A; Supplementary Table 1, Supplementary Video 1). Under our experimental settings (heat-killed bacteria), the *eat-2* longevity was shorter as previously reported on live bacteria but in line with previous findings that had demonstrated that *eat-2* predominantly extends lifespan by lowering the proportion of deaths caused by invasion of the pharynx from live bacteria (Zhao et al., 2019). Interestingly, our data showed that the longer-lived the mutant was, the more prolonged was the voluntary movement healthspan (Figure 1B). Therefore, interventions that increase lifespan also increase the time spent moving fast and actively.

**Figure 1:**
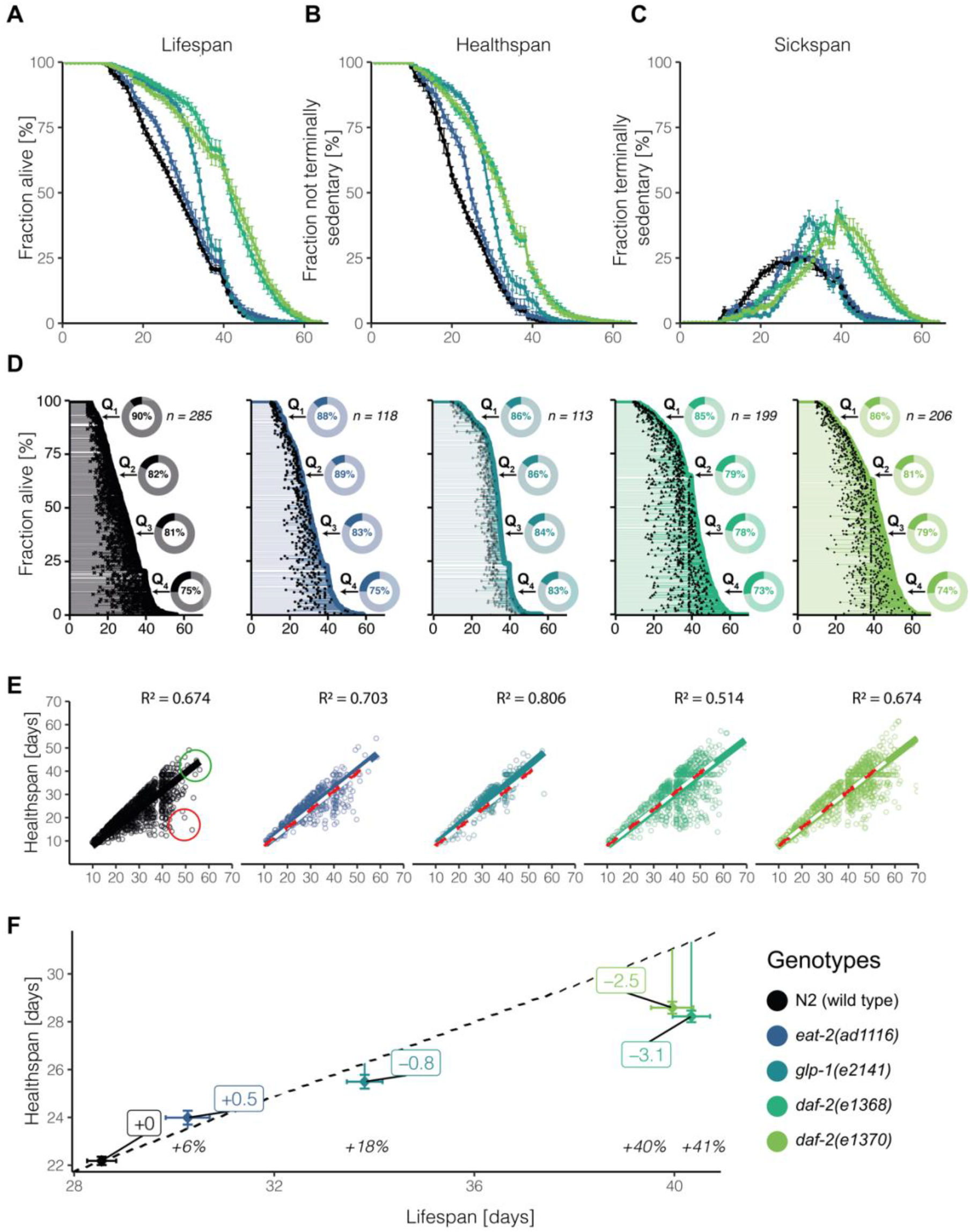
Voluntary movement quantification in aging *C. elegans* populations. *C. elegans* lifespan analysis displaying survival (A), healthspan (B), and sickspan (C) for each genotype. Error bars = S.E.M. between plates at 24h intervals. Healthspan refers to the timespan of fast movement, sickspan to the time spent in sedentary movement, and lifespan to the time until the animal fails to move irretrievably. The fate of each individual is displayed separately for each genotype overlayed with the population’s survival (D). Here, each individual’s healthspan is marked as a transparent line spanning from young adulthood to the onset of sickspan marked by a black dot and then extends further as sickspan until the individual’s point of death on the population survival curve. The inset displays the overall proportion each genotype spends in their healthspan for each lifespan quantile (Q_1_ – Q_4_). The correlation between health and lifespan is shown in figure (E). Each individual is represented as a point with its lifespan on the x-axis and its corresponding healthspan on the y-axis. A linear model passing through the origin is shown as a solid line. The wild type (N2) model is superimposed on the longevity mutants as a red and white dashed line. All genotypes are compared to temporarily scaled wild type N2 with the mean population life- and healthspan and error bars indicating the standard errors (F). The extrapolated ratio of health-to the lifespan of wild type (0.78) is displayed as a dashed black line. The distance of each population average is marked by a vertical line, and the difference in expected healthspan is indicated.

### Relative increase for both health- and sickspan in long-lived mutants

To better understand the rescaling of the time spent in frailty in these long-lived mutants, we analyzed the fraction of slow-moving animals per day. We observed gaussian activity distributions, which were shifted along the time axis for these longevity mutants (Figure 1C). This delayed onset of the sickspan (Figure 1C) is consistent with the prolonged healthspan of these long-lived mutants (Figure 1B). However, except for *eat-2* mutants, the width and the area of the Gaussian distributions were bigger for long-lived mutants than wild type (Figure 1C), suggesting an overall increase of sickspan. Thus, based on voluntary movement tracking, long-lived mutants display increased absolute health- and absolute sickspan compared to wild type. Next, we asked whether the fraction spent in health- and sickspan during the lifespan is altered. Wild-type animals spent 78% of their lifespan fast-moving and 22% slow-moving (Figure 1D). For long-lived mutants, we recorded about 70-79% of their lifespan are spent fast-moving (Figure 1D, Supplementary Figure 1), suggesting no compression of sickspan but rather a proportional scaling of both health- and sickspan relative to their lifespan.

### Heterogeneity in the length of sickspan but a fixed onset of sickspan

Since we did not observe a compression of voluntary movement sickspan of the entire population, we wondered whether individual animals that outlived their siblings would display a compressed sickspan. When we compared the sickspan traces of individual *C. elegans* for each genotype, we were surprised to measure such a vast heterogeneity (Figure 1D), given that all these individual animals of a population are genetically identical, consume the same food, and are housed in the same environment. Only the *glp-1*-mediated longevity showed an overall compression of individual sickspan traces (Figure 1D). For comparison among these different genotypes, we decided to use “relative age” by dividing lifespan curves into quartiles and computing the health-to-sickspan ratio for each quartile (Figure 1D insets; Supplementary Figure 2). Consistent with previous reports on wild type (Zhang et al., 2016), we found that in the first quantile of the lifespan curve, individual animals spent about 90% of their lifetime fast-moving and 10% slow-moving, indicating that these animals die young with a compressed sickspan compared to the last quartile wherein animals spent about 75% of their lifetime fast-moving and 25% slow-moving (Figure 1D insets; Supplementary Figure 1). Remarkably, it looks like the onset of an individual’s sickspan is a fixed event starting approximately when the first 10% of the isogenic population starts to die (Figure 1D). This observation suggests that up to a certain time point, the animal’s physiological integrity is maintained. After this time point, there appears to be a stochastic decay resulting in a heterogenous sickspan distribution. Viewing the data using this alternative interpretation of a fixed onset of sickspan would explain why animals in the first quartile of the lifespan curve die young and spend less time in poor health, while animals in the last quartile of the lifespan curve die old and spend more time in poor health. Thus, the time spent fast-vs. slow-moving seems to have a fixed onset in time.

### Voluntary movement healthspan temporally scales with lifespan except in *daf-2* mutants

The model of a fixed onset-timepoint for frailty would suggest that longevity interventions would simply delay the onset. To address this, we contrasted the number of days spent fast-moving (healthspan) for each individual as a function of their time lived (lifespan in days; Figure 1E). We found that the time lived correlated and predicted the time spent fast-moving with an R squared of 0.7 for wild type and R squared ranging from 0.5 to 0.8 for the longevity mutants (Figure 1E). Furthermore, the *glp-1* with an R squared of 0.8 and *daf-2(e1368)* with an R squared of 0.5 indicate lower or higher heterogeneity, respectively, compared to wild type (Figure 1E). This is also apparent in the increased or decreased spread of data points below the regression line in Figure 1E and by increased or decreased lengths of the individual sickspan traces in Figure 1D, respectively. One interesting aspect to note is that individuals in quartiles 2 and 3, which expire in the middle of the lifespan curves, displayed shortened healthspan relative to their lifespan, whereas individuals in the last quartile showed an extended healthspan relative to their lifespan (Figure 1E). This might be because sicker individuals simply died earlier, leading to an enrichment of healthier-aging individuals in the last quartile (Supplementary Figures 1, 2). Based on the high R squared values for all genotypes, we applied a linear model to investigate the relationship between health- and lifespan (Figure 1E). Steeper linear regression lines compared to wild type would indicate an increase in health- to lifespan ratio. The slopes of the linear model were steeper for *eat-2* and *glp-1*, but less steep for the two *daf-2* mutants compared to wild type (Figure 1E), suggesting that *glp-1* and *eat-2* spent a larger fraction and *daf-2* mutants spent a smaller fraction of their lifespan actively moving. Since slopes of linear models can be sensitive to extreme values, we compared the population means of health- and lifespan across all genotypes (Figure 1F). When we extrapolated the mean healthspan to mean lifespan ratio of wild type, we found that *eat-2* and *glp-1* were close to this extrapolated line, whereas the *daf-2* mutants lacked approximately three days (*i*.*e*., 7%) of mean healthspan in respect to their mean relative lifespan (Figure 1F). To demonstrate that all these measurements are true under other experimental settings, we chose temperature-sensitive sterile mutant *spe-9(hc88)* to compare to *glp-1(e2141)* that were raised at 25°C until day two of adulthood to avoid progeny and then kept for the remainder of their lifespan at 20°C and quantified comparable results (Supplementary Figure 3). Thus, we uncovered that the prolonged voluntary movement healthspan temporally scales with the prolonged lifespan for each of these longevity mutants except less stringently for *daf-2* mutants.

### Acoustophoretic characterization of *C. elegans* force and muscle power

Thus far, our observations and interpretations on healthspan are based on the decline of voluntary movement on culturing plates in the abundance of food. Certain genotypes like *daf-2(e1370)* are less motivated to forage and display a more rapid decline in voluntary foraging behavior compared to wild type leading to the interpretation of being less healthy (Hahm et al., 2015). In our setting, this lower foraging behavior is less pronounced in *daf-2(e1370)* since they were cultured at 15°C, an environment that avoids improper dauer program activation (Ewald et al., 2018). To overcome this, led us to develop an inducible, motivation-independent exercise platform for *C. elegans*. Our goal is to address the following shortcomings of current methods: the movement should be inducible with a strong stimulus and not dependent on secondary cues like food or intrinsic motivation, it should be measurable in a short time window to assess health in this instant, and it should directly measure a physiologically relevant parameter like maximum muscle force or functional tissue integrity. This is especially important when comparing different genotypes, which often respond differently to their environment.

In humans, one of the best predictors for all-cause mortality is the decline in muscle maximum force and power (Kostka, 2005; Leong et al., 2015; Petrella et al., 2005). However, a tool or device to quantify the maximum force and power of *C. elegans* muscles did not exist. The application potential would be immense since *C. elegans* muscle structures are strongly conserved, as in mammals, and forced maximum strength measurements to the point of collapse would be unethical in mammalian models. We developed a microfluidic device harnessing the power of acoustic standing waves (Figure 2A, 2B, Supplementary Video 2, for details, see *Cell Reports Methods*). We have recently applied ultrasonic waves to compress, move and quantitatively characterize larval *C. elegans* (Baasch et al., 2018). We reasoned that we could employ ultrasonic standing waves to trap and stretch out *C. elegans* in the minima of the acoustic force fields (Figure 2C, 2D). *C. elegans* dislike being trapped and try to escape by applying mechanical forces (body bending) against the acoustic force field (Figure 2C, 2D). The further away from the acoustic force field minimum, the harder it gets to move against the force field (Figure 2D). Suppose the animal is stronger than the applied acoustic force field. In that case, it can turn around in the microfluidic chamber (Figure 2C), typical escaping behavior of *C. elegans* known as omega reversals (Donnelly et al., 2013). Thus, the degree of deflection of the *C. elegans* body away from the acoustic force field minimum provides an estimate of the maximal muscle strength the animal can master to try to escape the acoustic trap.

**Figure 2:**
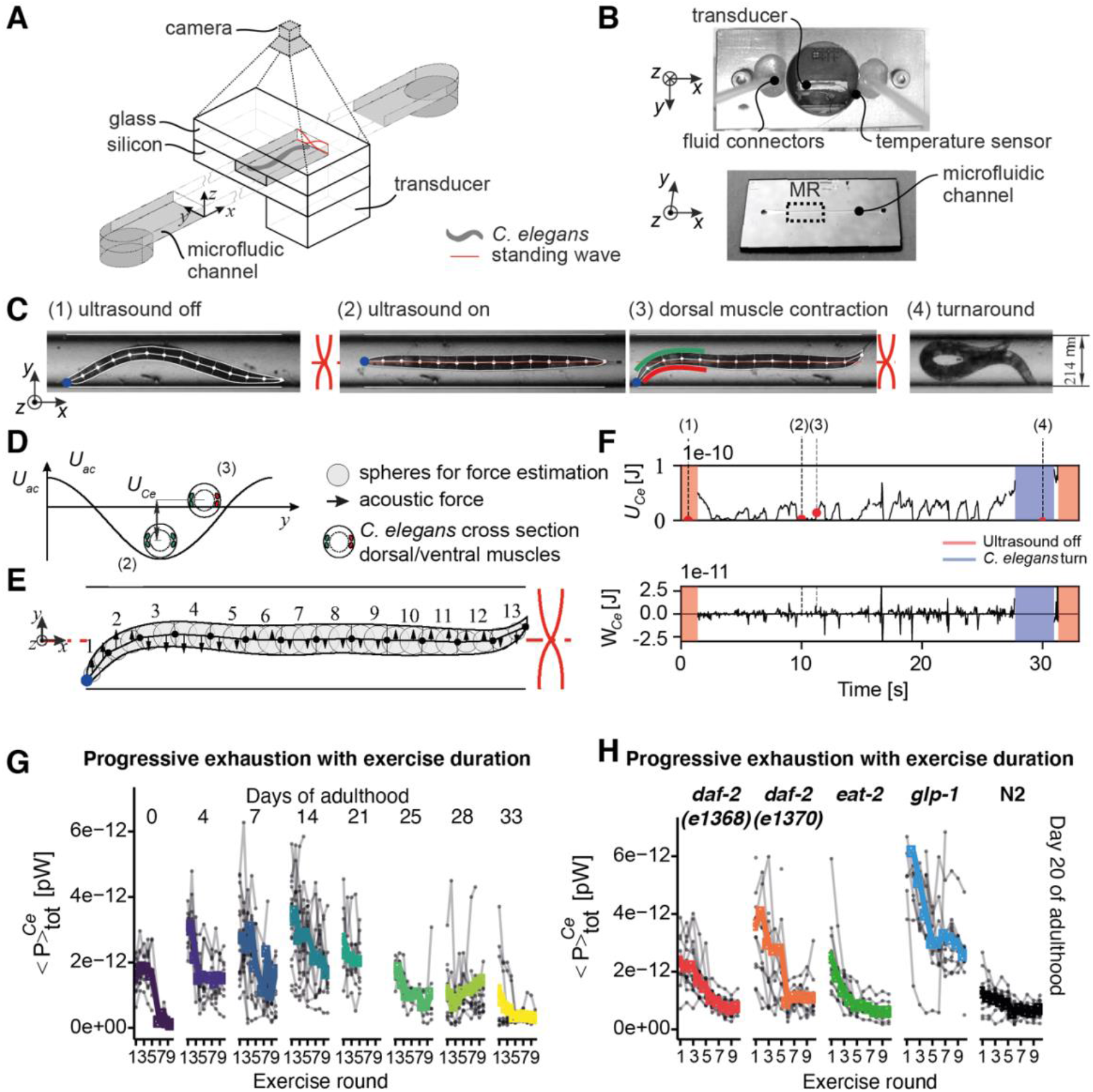
Experimental setup to measure maximum *C. elegans* muscle strength. Top view of the schematic representation of the silicon/glass acoustofluidic chip, fluid in- and outlets, and the piezoelectric transducer positioned at the bottom and *C. elegans* trapped in the standing wave (A). A photograph of the back of the chip together with the metal clamps, the attached fluid connectors, temperature probe, and a piezoelectric transducer is shown (B, top). The front view of the chip provides an overview of the device shape, the microfluidic channel, and the measurement region (MR) (B, bottom). A day 4 wild type *C. elegans*, as seen by the camera in the measurement region, is displayed together with the image processing output highlighting the outline and the segmented midline of the animal as well as the channel borders and centerline (C). The effect of the acoustic field (frequency: 3.543 MHz, voltage amplitude: 76 Vpp, 2 mode) is shown in (C, 2), aligning the animal at the midline. The animal exercises maximum muscle power attempting to bend its head away from the midline (C, 3) to achieve a turn (C, 4), likely as an attempted escape response. The working model to quantify the *C. elegans* muscular force consists of 13 rigid links along the animal’s midline, which are connected by joints. Blue arrows reflect the muscle activity acting on each joint to generate a force against the acoustic field (E). The acoustic force acting along the animal’s body is modeled as the individual acoustic forces acting on the grey spheres aligned along the body and represented by black arrows, with the length of each arrow being proportional to its force or moment magnitude. The acoustic radiation potential is illustrated using a *C. elegans* cross-section highlighting the animal’s four body wall muscles (D). If stretched, the animal rests at the minimum (see C 2) and, upon muscle contraction, moves upwards in its potential energy well (see C3). The four *C. elegans* frames (C 1-4) are put in the context of one 30 seconds actuation cycle showing the time-resolved total energy and mechanical work quantification for this animal (F). Red areas reflect regions of zero energy due to the acoustic field being turned off, and the blue areas indicate *C. elegans* turn movements during which muscle force estimation is paused. A C. elegans maximum force assessment routine consists of multiple 30 seconds exercise rounds intermitted by 5-second breaks for a total of up to ten actuation cycles. For wild type, this exercise regimen is displayed with the exercise rounds on the x-axis, the total power (time-averaged) on the y axis, as derived from the respective total energy curve in (F), and the age of the population given at the top of each facet (G). Wild type *C. elegans* is contrasted to long-lived *C. elegans* genotypes across up to ten exercise rounds focusing on the aged cohort above day 20 of adulthood (H).

### Muscular strength declines in aging *C. elegans*

To quantify muscular forces, we developed a model by dividing the *C. elegans* body plan into 13 rigid links connected by joints along the animal’s midline (Figure 2E). Upon applying acoustic force fields, we measured the deflection of these 13 nodes for 30 seconds and the number of times the animal escaped the force fields (Figure 2F). A typical exercise round is structured in up to ten cycles consisting of 30 seconds of ultrasonic force and a 5-second break (Figure 2G). We measured the muscular forces of aging wild-type *C. elegans* (Figure 2G). After 3-5 cycles, we observed muscle fatigue, which set in earlier the older the animals were (Figure 2G). We observed first an increase and then a decrease in the heterogeneity of individual *C. elegans* muscular strengths (Figure 2G). Similarly, we first saw an increase and then a progressive decline in muscle power during aging (Figure 2G). By contrast, we found that longevity mutants performed better in terms of muscle strength and function at day 20 of adulthood (Figure 2H). This indicates the preservation of muscle power in aging longevity mutants.

### Longevity mutants showed prolonged healthspan assessed by the strength performance in longitudinal comparison to wild type

The overall force and power of a *C. elegans* depend on its muscle strength as well as its total body size. In agreement with previous reports (Hulme et al., 2010; Shi and Murphy, 2014), we observed adult *C. elegans* kept growing in body size beyond the reproductive period and then shrunk during aging (Figure 3A, 3B). A prolonged growing phase correlates with longevity (Hulme et al., 2010). We found that longevity mutants prolonged their growing phase and shrunk less than wild-type animals during aging (Figure 3A, 3B). Structural integrity declines during *C. elegans* aging, such as internal organ atrophy (Ezcurra et al., 2018), loss of internal pressure (Gilpin et al., 2015), and disorganization of the exoskeleton cuticle (Essmann et al., 2020). We noticed that in the acoustic force field, *C. elegans* undergoes compression, and this compressibility stays fairly constant during aging (Figure 3C, 3D). We conclude that although morphological changes occur during aging, the mechanical properties regarding compression are less affected by age. This points towards muscular strength playing a pivotal role. On average, young *C. elegans* can overcome the acoustic force field leading to an omega turn eight times each cycle (Figure 3E, Supplementary Video 3). The ability to overcome the force field and turn in the microfluidic chip progressively declines during aging but is preserved in longevity mutants (Figure 3E, Supplementary Figure 4). Turning in the chip can be viewed as a measure of high-intensity muscular capacity since the animal can completely overcome the force field. It also showcases that the animal is not placid but trying to escape. Next, we assessed the overall energy per individual *C. elegans* as an assessment of overall body volume deflected against the force field. We found an increase of energy per individual until mid-age and then a decline (Figure 3F) reminiscent of the longitudinal body size curve (Figure 3G). In our measurements to determine the overall force and power of *C. elegans*, the body size is a confounding factor. In all our longitudinal measurements, we had included a positive control in the form of a muscle-defective mutant (CB190) that carries a mutation in the muscle myosin class II heavy chain (*unc-54*). These muscle-constriction-defective mutants were unable to perform omega turns in the chip but showed similar compressibility and longitudinal growth curves, illustrating that the rise and fall of the overall energy are confounded by the organismal growth curve. Therefore, we decided to use the dynamic power as defined as energy expenditure relative to the previous time point and normalized it by the volume of each animal (*i*.*e*., volume; Figure 3H). Using a human analogy, total refers to how long a weight can be lifted, dynamic power only considers the process of lifting the weight without holding the weight, and dynamic power takes the weight of the person lifting the weight into account. In this way, we found that the overall force and power of longevity mutants were preserved for almost three-quarters (55-83%) compared to about one-third (30%) of their lifespan in wild type (Figure 3I). Thus, muscular strength is maintained longer in longevity mutants.

**Figure 3.**
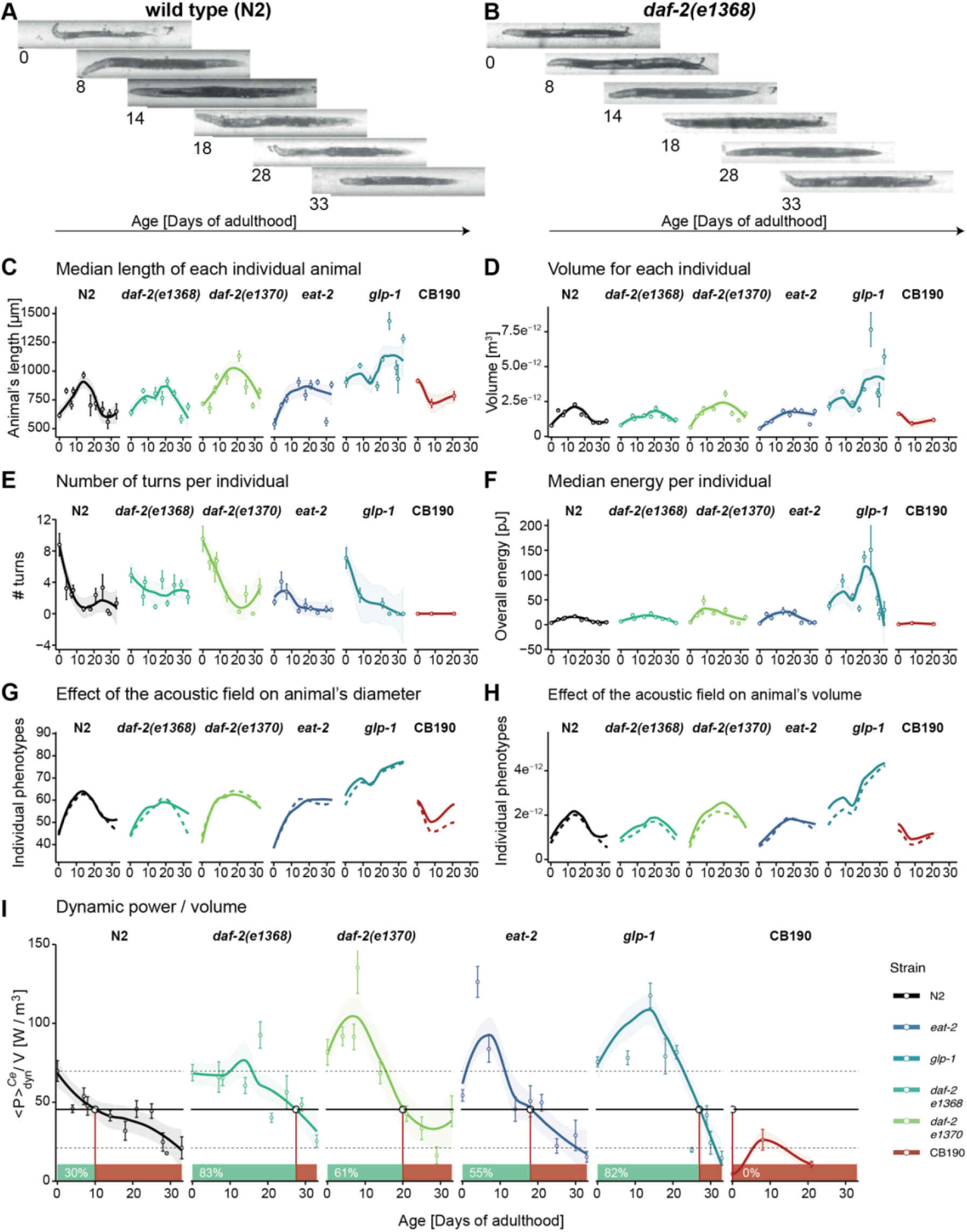
Time course of fitness- and structural-related phenotypes for wild type and long-lived *C. elegans* under acoustic stimulation. Acoustic compression was used to quantify phenotypes directly (# turns, energy, diameter- and volume compression, and dynamic power by volume) as well as indirectly by exploiting the non-destructive linear alignment of the animals (length, volume). Randomly selected images for N2 (A) and *daf-2(e1368)* (B) at different ages are displayed to illustrate the positioning of the *C. elegans* in the microfluidic measurement channel and how all matrices are shown below were obtained. Measurements are displayed as mean ± standard error for each assessed age point and subjected to local polynomial regression fitting displayed as a full line with the confidence interval set to 95% and bounded by dashed lines. For the acoustic compression only, the fitted line is shown. The length (C) and volume (D) of the animal was quantified automatically using the entire length and area of the animal in the channel, respectively. The number of turns was quantified manually and corresponded to the number of times an animal successfully changed its orientation in the channel by 180°. The total energy of the individual was calculated using the magnitude of the lateral deflection of the animal from the channel middle (F). The compression experienced by the animal diameter (G) and volume (H) when the field is activated was computed automatically and displayed as a full line when the field is on and a dashed line when the field is off. The dynamic power/volume (I) reflects the work the animal performs against the field to change its lateral position in the channel and is normalized by the overall volume of the animal to enable comparisons across genotypes making this the most informative health parameter. The mean value for day 0 and day 33 N2 animals are indicated in black dashed lines, and the half-activity value between these two extremes is shown as a full line which is also used to deduce the health- to sickspan transition for each genotype. Using this N2 half-activity value, the individual strains reach their sickspan at approximately 10 days for N2, 28 days for *daf-2(e1368)*, 20 days for *daf-2(e1370)*, 18 days for *eat-2(ad1116)*, and 27 days for *glp-1(e2141)* while CB190 spends its entire lifespan in its sickspan fraction. Long-lived genotypes, in general, do not experience the same linear energy density decline as wild type since their total energy decrease is slowed down, as is their growth leading to a non-linear energy density trajectory.

### Temporal scaling of age-related pathologies in longevity mutants

Next, we asked whether other age-related pathologies or morphological changes show any delayed onset in longevity mutants compared to wild type. We quantified 592 animals, investigated timepoints between day 0 and day 33 (12 animals on average per strain and time point) at the first two cycles of actuation (1183-time sequences), which comprised over 800’000 frames in total. We then manually quantified additional morphological changes such as intestine length and diameter, pixel intensity, wrinkles in the cuticle, and pharynx diameter in a subsampled representative subset (approx. 50’000 frames; Supplementary Figure 5). Although not all, many age-related phenotypes were delayed in their onset and displayed a slowed decline in longevity mutants compared to wild type (Figure 4A, 4B, Supplementary Figure 6). Rescaling phenotypic trajectories of wild type by the lifespan extension observed in the long-lived strains revealed that many closely match the trajectories observed in long-lived strains for both *daf-2* mutants (Figure 4C). Notably, for animals’ length, diameter, volume, and intestine length, phenotypic scaling was observed when comparing wild type to *daf-2(e1368)* and *daf-2(e1370)*. In the case of *eat-2*, the observed lifespan extension was too limited to draw conclusions, and *glp-1* never ceased growing. The severely paralyzed myosin mutant *unc-54(e190)* displayed the opposite phenotypic trajectories than all other genotypes (Figure 4C). When approximating the phenotypic trajectories as segmented linear fit reflecting the separated phases of growth and decline, we observed that the starting values as young adults are often similar (Figure 4C). However, the slopes and point of decline are shifted compared to wild type. Investigating each phenotype in isolation is hindered by the inherent noise in the measurement as well as by the incomplete picture each phenotype provides. Furthermore, many phenotypes like length, diameter, and volume were strongly correlated. For this reason, we subjected all phenotypes to Principle Component Analysis (PCA) to study the overall age trajectory (Supplementary Figure 7A). We traced these phenotypes of all genotypes across the PCA plot as they age (Supplementary Figure 7B). All physiological parameters increased from young to middle-aged and then reverted again as the animal reached old age. However, muscle strength density decreased steadily. Using the paralyzed mutant, we were able to establish the bottom left quadrant as a reduced health area. This was only possible when using both physiological and performance measurements and was entered only by the paralyzed strain as well as old wild-type animals. The multi-phenotype traces are also shown for each genotype individually (Supplementary Figure 7C, 7D). Taken together, this suggested that many of these phenotypes change similarly during aging, that many are temporally scaled in longevity interventions, and that maximum muscle strength offers an orthogonal perspective on studying aging compared to physiological features. This highlights the importance of performing high-intensity muscle strength measurements when studying physiological aging and quantifying healthspan.

**Figure 4.**
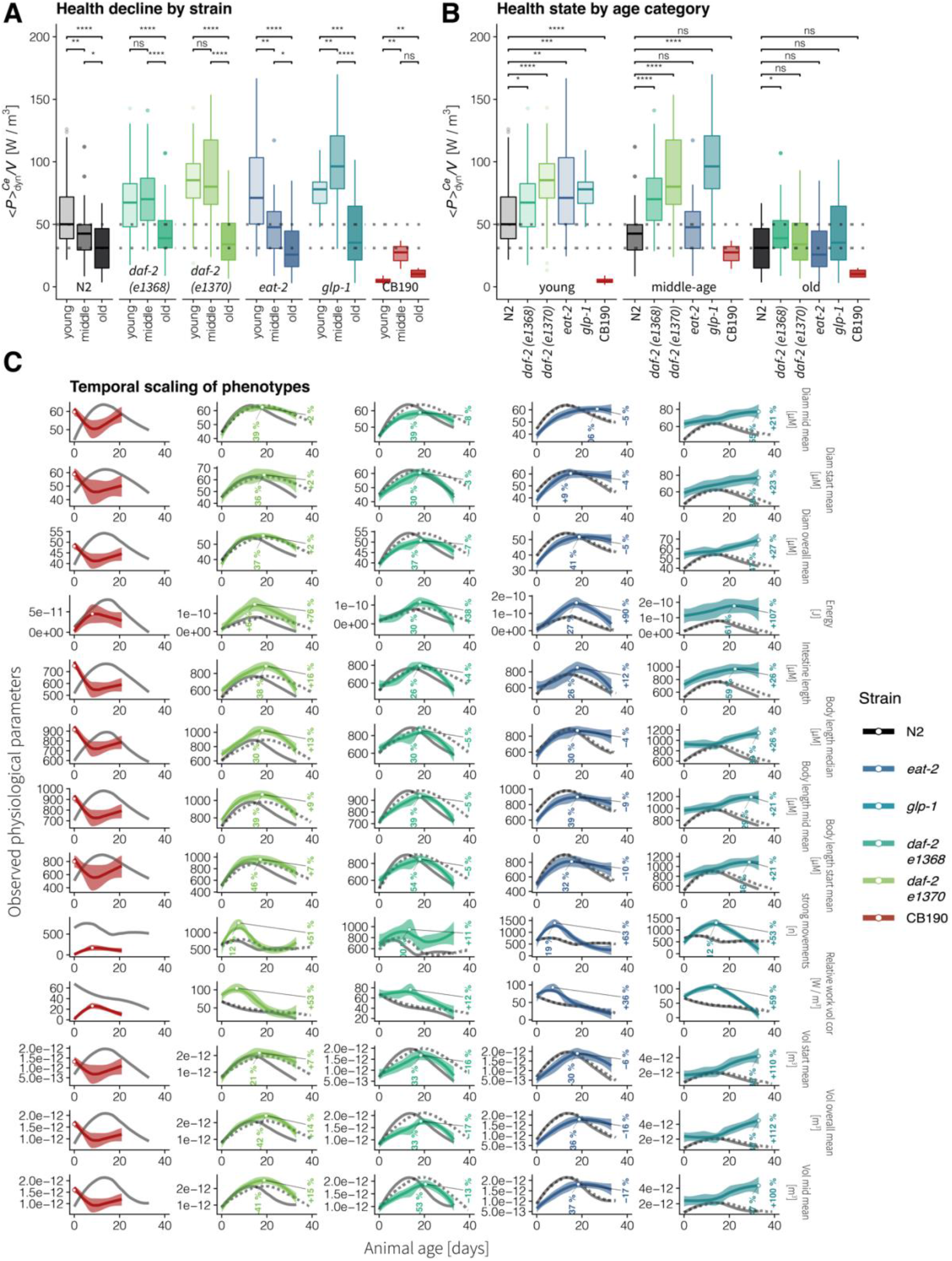
Temporal scaling of *C. elegans* aging phenotypes. The measured values of selected phenotypes are shown using different scales on the y-axis, animal age on the x-axis, and facetted by strain (A). To address the temporal scaling of phenotypes hypothesis, the loess fit of the measured values for wild type is shown as a solid grey line, and it is temporally scaled values using the respective mean lifespan increase experienced by the respective strain are shown as a dashed line also in grey. The measured phenotype trajectory for each strain is shown in color, each in a separate panel. The maximum fitted value is marked using. White point and its increase relative to the maximum fitted values measured for wild type are shown for both the age and phenotype variables. The same phenotypes as in subfigure (C) are modeled using a piecewise linear relationship in (B). The breakpoint of the segmented fit is estimated by the model at the age value, where the linear relationship between the measured phenotype and population age changes. The individual animals measured at each time point are displayed as mean +/- standard error.

### Longevity mutants show prolonged absolute but not relative healthspan

Our data revealed that longevity mutants stay healthier compared to wild type and experience a slower decline in physiological integrity. Indeed, dividing the lifespan of each genotype into three chronological fixed age categories: young (less than 7 days), middle (older than 8 but younger than 19 days), and old age (>20 days of adulthood), showed a progressing decline of volume-corrected work performed (Figure 5A, 5B). Longevity mutants performed better in the middle age group than wild type, but only *daf-2(e1368)* outperformed wild type in the old age group (Figure 5A, 5B). Using hierarchical clustering of temporally scaled phenotypes as a complementary analysis, we found that longevity mutants often cluster with chronologically younger wild-type samples (Supplementary Figure 8).

**Figure 5.**
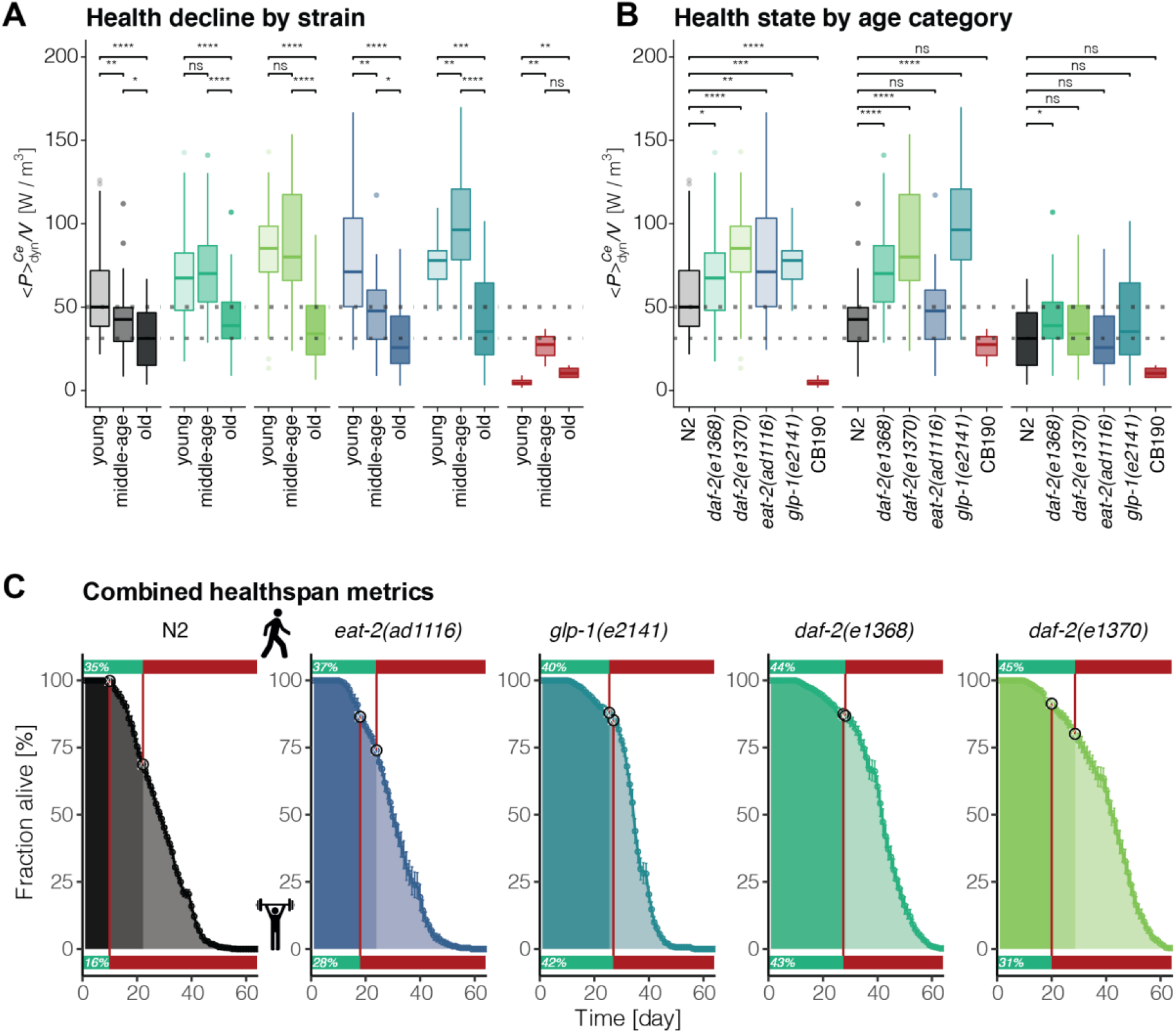
Integration of voluntary movement and forced maximum muscle strength quantification to yield a comprehensive understanding of *C. elegans* healthspan. The volume-corrected relative work performed by each genotype and grouped by age category (young <= 7 days, 8 < middle < 20 days, old >= 20 days) is shown as boxplots facetted by genotype (A) and by age category (B). P-values of selected comparisons (Mann-Whitney test) are displayed as symbols (ns > 0.05, ^*^ < 0.05, ^**^ < 0.01, ^***^ < 0.001, ^****^ <0.0001). The Population medians for young and old N2 are displayed as horizontal dashed lines across all panels. C) The measured lifespan of each *C. elegans* genotype is displayed with animal age on the x-axis, the fraction of the population that is alive on the y-axis, and the mean and standard error between plates shown as point and line range. The voluntary movement was quantified using the active vs sedentary behavior of the unstimulated animals on plates and is shown for each population as a bar at the top of each panel. Muscle health and the corresponding onset of sickspan due to reduced muscle function is depicted as a bar at the bottom of each facet. The position on the lifespan curve corresponding to the health-to-sickspan transition of either the unstimulated or stimulated healthspan quantification is marked by a white circle. The two health assessments divide the lifespan curve into three segments, with decreasing health status reflected by increasing transparency.

### Integration of voluntary movement and forced maximum muscle strength quantification to yield a comprehensive understanding of *C. elegans* healthspan

Having two independent assessments of healthspan that act on very different stringency levels, we can further divide *C. elegans* healthspan into 3 divisions: prime health (passes both matrices), fragile health (passes 1 metric), and sickspan (failing both matrices) (Figure 5C). Consistent with previous observations in other species, the maximum power drops prior to the cessation of general mobility (Kostka, 2005; Leong et al., 2015; Petrella et al., 2005). Muscle performance is much more improved in longevity mutants in relation to wild type compared to the voluntary movement (Figure 5C). Integrating both matrices revealed that wild type spent around 16% of their lifespan in prime health, whereas longevity mutants spent double the time in prime health (28-42%; Figure 5C). From all four longevity genotypes, *daf-2(e1368)* appears to be the healthiest strain (Figure 5C). Thus, combining multiple matrices of physiological and behavioral integrity is a powerful assessment of healthspan.

## Discussion

Understanding the relationship between healthspan and lifespan is an important question in aging research since geroscience aims to increase the time spent in good health and to postpone and compress the time suffering from age-related pathologies and chronic diseases (Kaeberlein, 2017; Kennedy et al., 2014; Olshansky and Carnes, 2019; Partridge et al., 2018). Model organisms like *C. elegans* are used to identify longevity-promoting interventions that can then be of translational value for humans (Kenyon, 2010; Magalhães et al., 2017). There is a fierce debate whether *C. elegans* longevity interventions show compression of sickspan and are of translational value for improving healthspan or healthy aging in humans (Bansal et al., 2015; Hahm et al., 2015; Huang et al., 2004; Podshivalova and Kerr, 2017; Stamper et al., 2018). In this study, we set out to develop a robust method to quantify maximum muscle strength as a highly interpretable healthspan metric with translational value. Lifespan measurements are well-established and allowed the study of hundreds of lifespan-extending compounds and genetic alterations, leading to ground-breaking discoveries. However, this development is not reflected in the area of healthspan extension. Numerous methods exist which often measure proxy phenotypes for healthspan that are also motivation dependent like pharyngeal pumping, thrashing, and others. With our approach, we were able to directly quantify muscle health in *C. elegans*. This is especially relevant since *C. elegans* is an ideal model system for large-scale genetic screening, and both the microfluidic device as well as the image detection can be multiplexed. This approach could translate to the much-needed identification of (muscle) health-promoting interventions.

The microfluidic device operates using acoustophoresis to generate an acoustic force field to quantify the physical fitness and muscle strength of aging *C. elegans*. Using different ways to assess healthspan in forms of voluntary movements, muscular force, muscular fatigue, structural integrity/compressibility, and quantifying several age-related morphological changes, including cuticle/skin wrinkles, body, and internal organ sizes, we find that most of these phenotypic changes are postponed in longevity mutants. We observed a hierarchical and time-dependent succession in the occurrence of these phenotypes, starting with a decline in maximal force as indicated by overcoming acoustic field (omega turns), a decline in dynamic power, seizing of body and organ growth (intestine), and then decline in voluntary movement and becoming inactive and lethargic. Using principle component analysis, we show that many of these phenotypes are strongly cross-correlated. The delay of all these age-related phenotypic changes is evident when using chronological age as a reference point for comparison of longevity mutants with wild type but disappears when using relative age as a reference. This points to the idea of temporal scaling of the healthspan. Consistent with this idea, we find that sickspan is neither compressed nor prolonged in longevity mutants compared to wild type. Thus, our quantifications suggest that *C. elegans* healthspan undergoes temporal scaling in longevity.

Aging is defined as a set of phenotypes or senescent pathologies occurring with a higher proportion in older individuals (Freund, 2019). Which senescent pathologies limit the lifespan depends on the context and are different for different species, genotypes, and environments (Freund, 2019; Gems, 2015). Whether our chosen set of phenotypes assessed are directly limiting or affecting lifespan is unclear. However, it is evident that not one single mechanism underlies all our measured age-related phenotypes. On the other extreme, we do not observe a “one mechanism causing one age-related pathology” mechanism. Our data shows that some phenotypes correlate and also follow a hierarchal time-dependent order of occurrence, indicating that these senescent pathologies are interconnected. This favors a mixed model of several causal mechanisms affecting multiply connected and independent senescent pathologies/phenotypes, including lifespan limiting phenotypes (Freund, 2019; Gems, 2015). Even if we might not measure lifespan limiting phenotypes directly, the interconnectedness of phenotypes should reveal the same picture of temporal scaling of age-related pathologies.

The lifespan of *C. elegans* can be increased by up to ten-fold (Ayyadevara et al., 2008) and decreased by 40-fold, but surprisingly the lifespan curves often follow the same rescaled distribution (Stroustrup et al., 2016). Temporal scaling was also noted when comparing expression profiles of longevity mutants compared to wild-type *C. elegans* (Tarkhov et al., 2019). Temporal scaling might also be the underlying reason why “aging clocks” based on transcriptional profiling work and longevity mutants’ biological age determined by the clock is younger than their chronological age (Meyer and Schumacher, 2021). Furthermore, temporal scaling was also observed for bacterial aging (Yang et al., 2019), suggesting that temporal scaling is an ancient underlying process conserved through evolution. Whether temporal scaling also occurs in mammalian longevity needs to be determined in the future. However, study designs for primary outcome measures for clinical trials on aging are based on the underlying assumption of temporal scaling. For instance, the Metformin in Longevity Study (MILES; NCT02432287) used RNA-sequencing of muscle and fat tissue to determine a rejuvenation to a younger expression profile as a primary outcome measure. Interestingly, certain longitudinal *C. elegans* phenotypes are comparable to human age-related phenotypes. Analogous to the *C. elegans* volume increase to peak mid-age and then decrease is that human BMI and waist circumference also follows this early-to-mid-life increase reaching a peak around 65-70 years and then declining (Kuo et al., 2020). Furthermore, grip strength progressively declines after the age of 30-40 years (Kuo et al., 2020), similar to *C. elegans* muscular strength. This raises the question of whether non-compression of sickspan observed in *C. elegans* means or interpolates to non-compression of sickspan in humans? Since aging is universal, it is tempting to speculate that the underlying mechanisms of aging or age-dependent phenotypes are also universal. This might be a potentially erroneous or unproven extension of the observation that almost all living things age (Freund, 2019). Although phenotypic changes, such as the loss of *C. elegans* muscle force, is analogous to loss of grip strength or muscle strength loss in humans, the underlying biological mechanisms resulting in physical weakness might be different. Our study makes no conclusion or interpolation about the compression of the sickspan in humans.

There is an accumulating body of evidence that long-lived humans are healthy during old age. For instance, 56-83% and 15-23% of centenarians, people over the age of hundred years, delay the onset of chronic age-dependent diseases and physical disabilities or were even free of such co-morbidities and frailties, respectively (Ailshire et al., 2015; Evert et al., 2003). Centenarians have lower incidence rates of chronic illnesses compared to their 90- or 80-year old matched-controls (Andersen et al., 2012; Evans et al., 2014; Ismail et al., 2016; Kheirbek et al., 2017). This also extends to family members related to centenarians compared to families without centenarians (Ash et al., 2015; Sebastiani et al., 2013). Thus, centenarians have a later onset and a lower rate of incidence compared to people in their eighties, similar to our observation when comparing longevity mutants to wild-type *C. elegans*. However, since centenarians get to enjoy at least 20 more years, how would this comparison look if we were to compare relative age to chronological age? Is sickspan compressed or temporally scaled in centenarians compared to the average population? As our life expectancy doubled in the last hundred years and we are on the course of potentially reaching the limit of our lifespan (Olshansky and Carnes, 2019), the accompanied delayed onset of disabilities already started to decelerate in longer-lived women compared to men (Freedman et al., 2016). On the other hand, there are several interventions that increase healthspan without increasing lifespan per se identified in mice (Fischer et al., 2016; Garcia-Valles et al., 2013) and Rhesus monkeys (Mattison et al., 2012). Thus, studying longevity is an important first step of identifying molecular mechanisms promoting healthy aging, but our study and others (Fischer et al., 2016; Garcia-Valles et al., 2013; Mattison et al., 2012) point toward that it is crucially important for geroscience to start investigating interventions that improve healthspan directly in future studies (Olshansky, 2018). Initial steps in defining healthspan (Kaeberlein, 2017; Kennedy et al., 2014) and also tools and experimental setups, including this study, are being developed to reliably quantify healthspan (Bellantuono et al., 2020; Collins et al., 2008; Haefke and Ewald, 2020; Teuscher and Ewald, 2018; Teuscher et al., 2019).

In summary, we have demonstrated that *C. elegans* sickspan is neither compressed nor extended in longevity mutants providing an alternative answer to an ongoing debate in the aging field. With our measurements, we showed that previous claims that insulin/IGF-1 receptor mutants have increased sickspan compared to wild type are correct if the voluntary movement is measured, but not the case if the muscular function or other healthspan measurements are considered that do not rely on the behavioral state of the animal. By adjusting the reference system from chronological age to relative age, we provide evidence that the healthspan of longevity mutants undergoes temporal scaling. Future studies using our acoustophoresis approach to study the role of healthspan will reveal novel strategies to improve healthy aging.

## Author contributions

All authors participated in analyzing and interpreting the data and designing the experiments. CS performed lifespan assays. PR designed and performed microfluidic assays. CS and PR analyzed the datasets jointly. CS and CYE wrote the manuscript in consultation with the other authors.

## Author Information

The authors have no competing interests to declare. Correspondence should be addressed to C.

Y. E. and J.D.

## Acknowledgment

We thank the Ewald lab for constructive comments on the manuscript. Some strains were provided by the CGC, which is funded by NIH Office of Research Infrastructure Programs (P40 OD010440). Funding from the Swiss National Science Foundation PP00P3_163898 to CYE and CS.

---

**Supplementary Figure 1:**
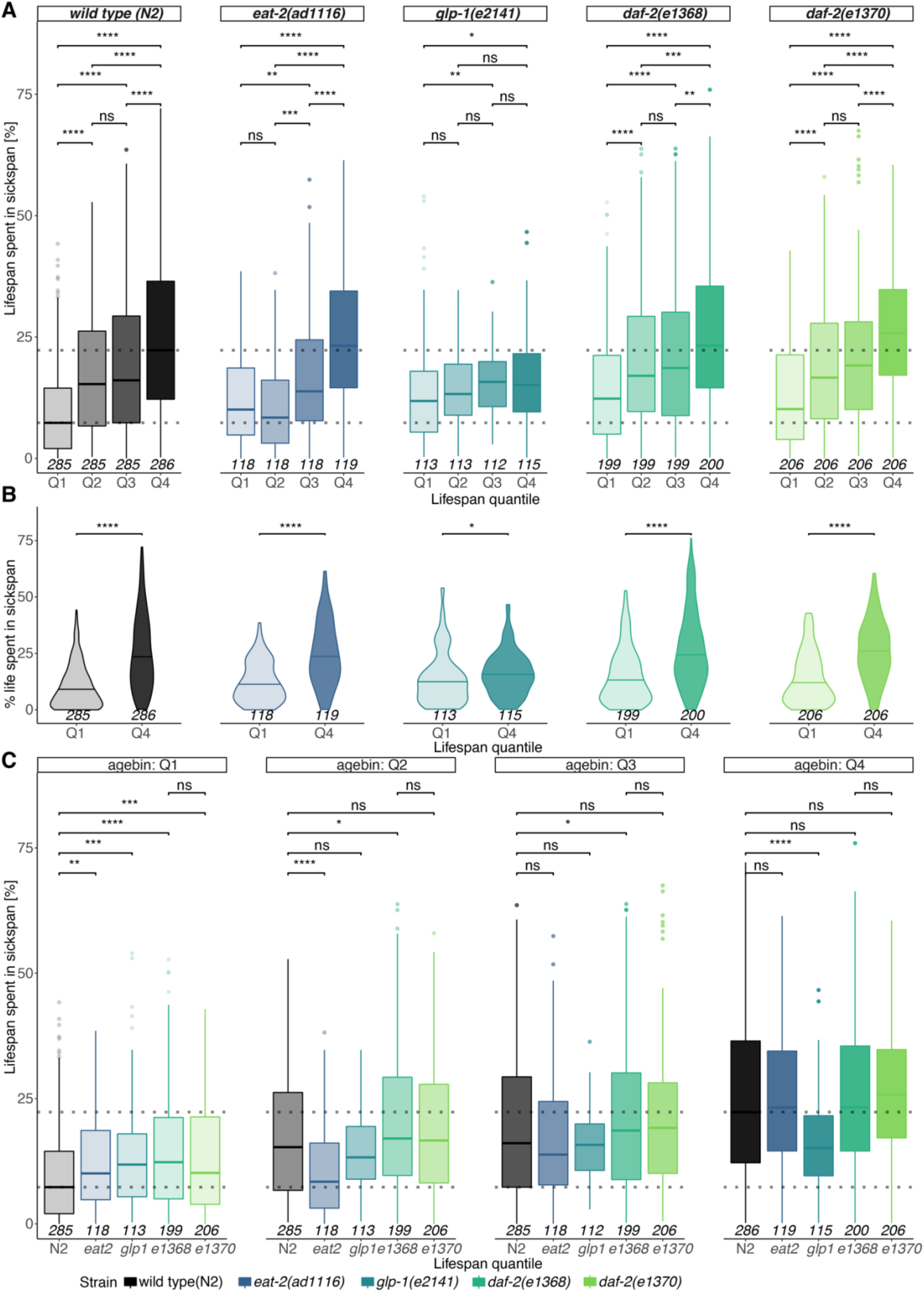
Paradoxon - young animals die apparently in good health. The sickspan distribution for every quartile of the lifespan distribution is displayed as boxplots, with each strain in a separate panel (A). Animals experience progressively higher sickspan ratios with each quantile, with the two middle quantiles being the most similar. The distributions of the first and fourth quartile are shown as violin plots with their median line highlighted by a horizontal segment showcasing the increase of relative sickspan and heterogeneity with age across genotypes (B). To compare the individual genotypes for each relative age cohort, the corresponding quartiles are contrasted as boxplots for each strain and each quartile in a separate panel and indicate temporal scaling of healthspan (C). The sickspan ratio is compared across genotypes and quartiles (Mann-Whitney test) and the P-values are depicted as symbols (ns > 0.05, ^*^ < 0.05, ^**^ < 0.01, ^***^ < 0.001, ^****^ <0.0001). When all quartiles are displayed, the median sickspan percentage of the first and last quartile of the wild-type population are indicated as dashed horizontal lines. The number of observations associated with every subpopulation is displayed at the bottom of each observation. Each strain is shown in its distinct color, and increasing age is reflected by increasingly dark coloring.

**Supplementary Figure 2.**
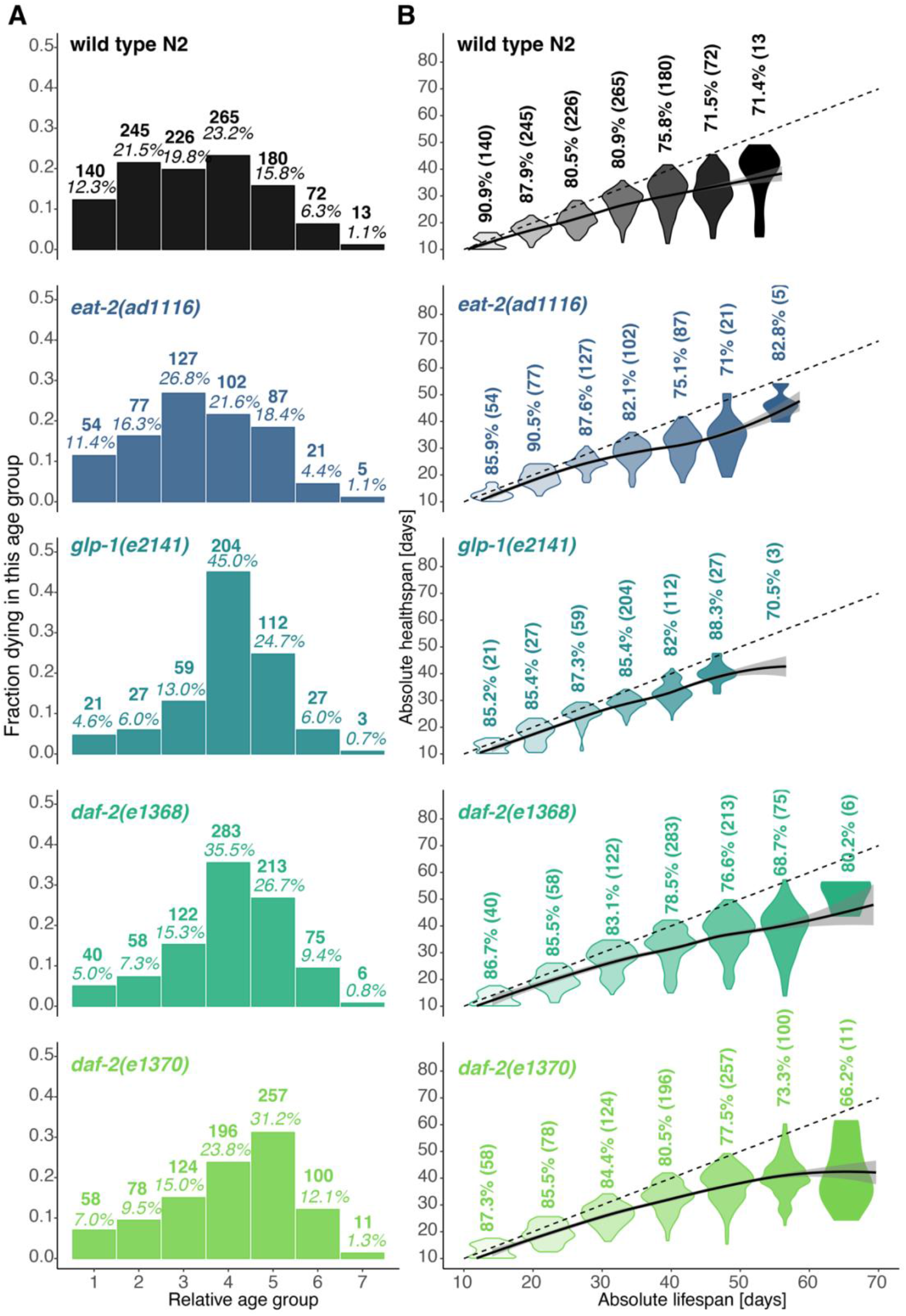
Relative healthspan within early and late dying *C. elegans* groups. To address the question of whether outliers drive the overall health-to-lifespan ratio, each genotype was divided into seven groups with equally sized age bins between the first and last recorded death in each genotype’s population to visualize the distribution of death events. At the top of each bar, the number of animals observed to die in this cohort and its share in the overall population is displayed **(A)**. The healthspan distribution in each age group is shown as a violin plot **(B)**. The average percentage of each population spent in their healthspan is displayed above each group, with the population size indicated in brackets. The health-to-lifespan diagonal is shown as a dashed line, and a loess trendline was fitted to the binned datasets with a 95% confidence interval shaded in grey. In case the sample size falls below a threshold, the violin plot was omitted. Each analysis is displayed separately for every *C. elegans* genotype, which are each shown in a separate line and in a distinct color. The spread of the healthspan distribution increases at older ages, with some animals experiencing nearly no sickspan and others spending most of their life in the sickspan portion.

**Supplementary Figure 3.**
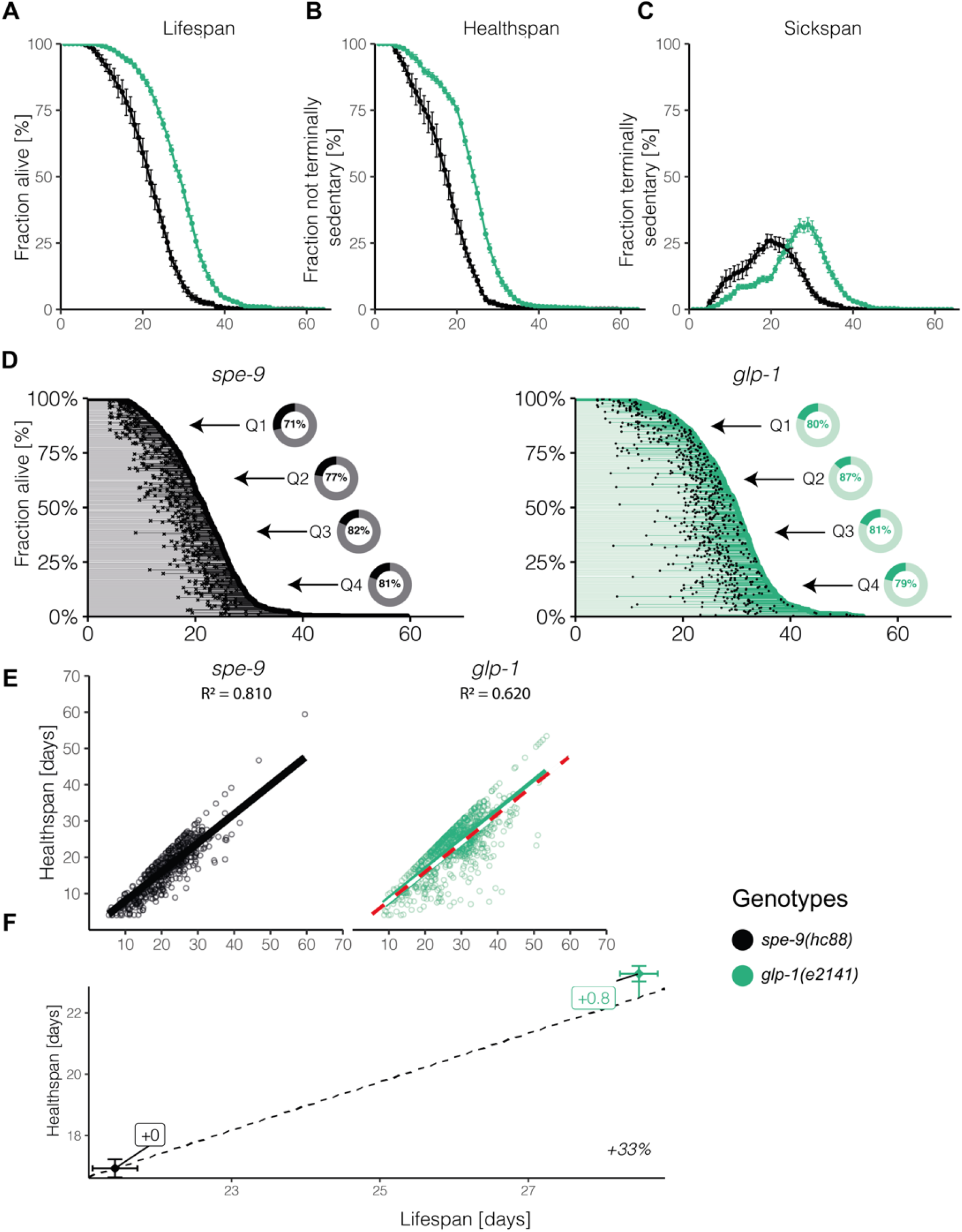
Voluntary movement quantification in aging *C. elegans* populations under different experimental setup conditions. To exclude any potential confounding factors due to experimental setup, we chose to compare temperature-sensitive sterile TJ1060 *spe-9(hc88)* as a normal-lived wild-type control to long-lived *glp-1(e2141)*. Both strains eggs were cultured at 25°C until day two of adulthood and then shifted to another bacterial food source (HT115, L4440) and maintained at 20°C for the remained of the lifespan in the lifespan machine. Shown is a composite of three independent trials. Comparable to Figure 1, (A) *C. elegans* lifespan analysis displaying survival, (B) healthspan, and (C) sickspan for each genotype. (D) Individual onset of sickspan marked by a black dot and then extends further as sickspan until the individual’s point of death on the population survival curve. (E) The correlation between health and lifespan is shown in figure. (F) The long-lived *glp-1(e2141)* is compared to temporarily scaled *spe-9(hc88)* with the mean population life- and healthspan and error bars indicating the standard errors. For details, see Figure 1 and Supplementary Table 1 for raw data and statistics.

**Supplementary Figure 4.**
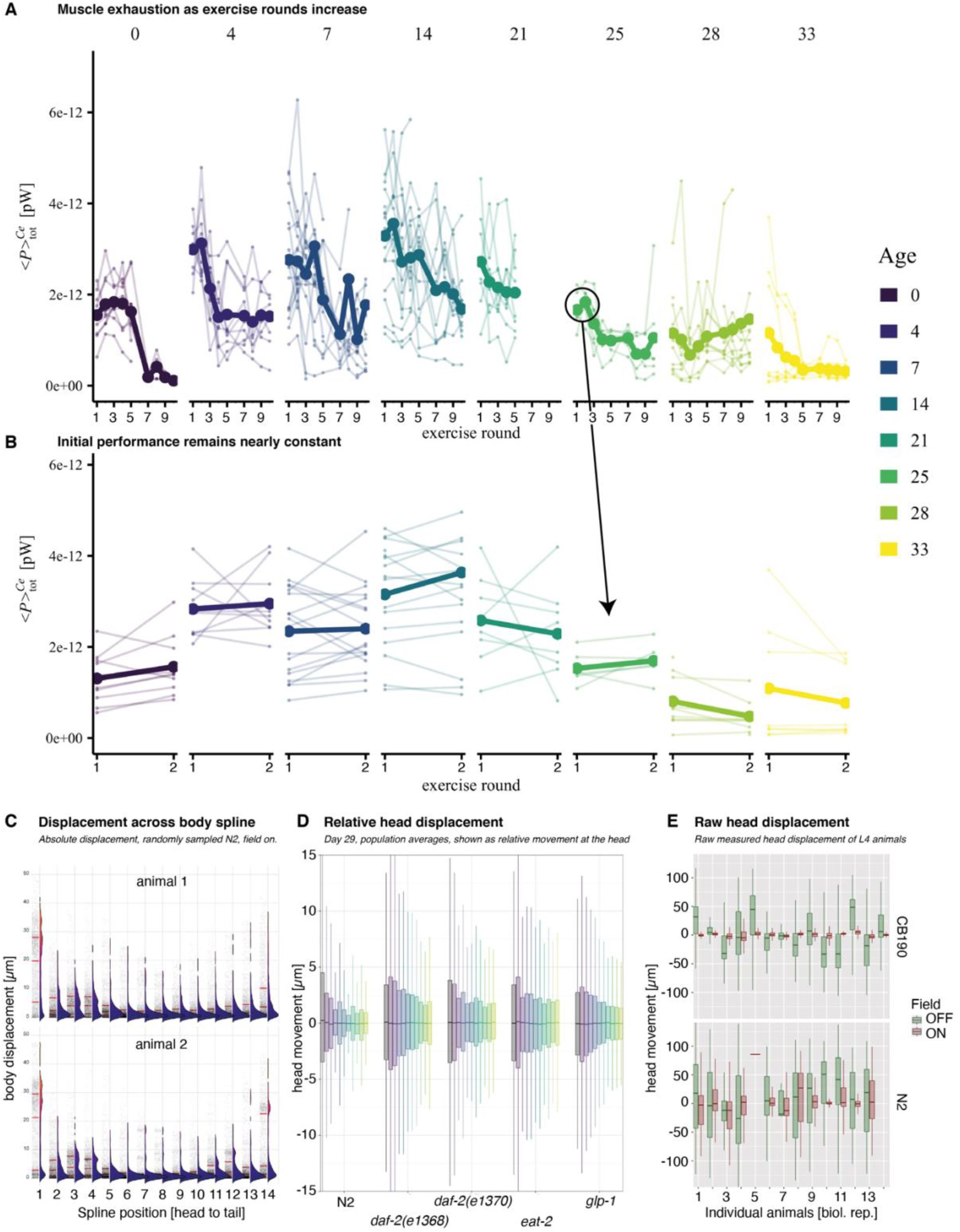
*C. elegans* movement in the acoustic exercise chamber. The exercise regimen was benchmarked for wild type (N2) *C. elegans* at different ages to quantify after how many cycles they experience muscle fatigue (A, B). Every trace corresponds to one individual animal that was up to ten times subjected to 30 seconds maximum force exercise followed by a 5 seconds break (A). Thicker lines represent the population means at every time point. To quantify maximum muscle strength, we selected the first two actuation cycles (B) at which no fatigue can yet be identified. The total energy is computed over the entire body and muscle apparatus of the animal. Interestingly, the spline position most activated in this exercise is the head and tail sections (C). Focusing on the head, we quantified the relative head movement for different strains at old age and also observed the same trend of decreasing movement with higher actuation cycles (D). The extreme case of the paralyzed CB190 vs. N2 in young adulthood displays that while the mutant is still able to move the head away from the midline of the field when the acoustic field is off, it is unable to move when the field is on while N2 successfully fights against the applied forces (E).

**Supplementary Figure 5.**
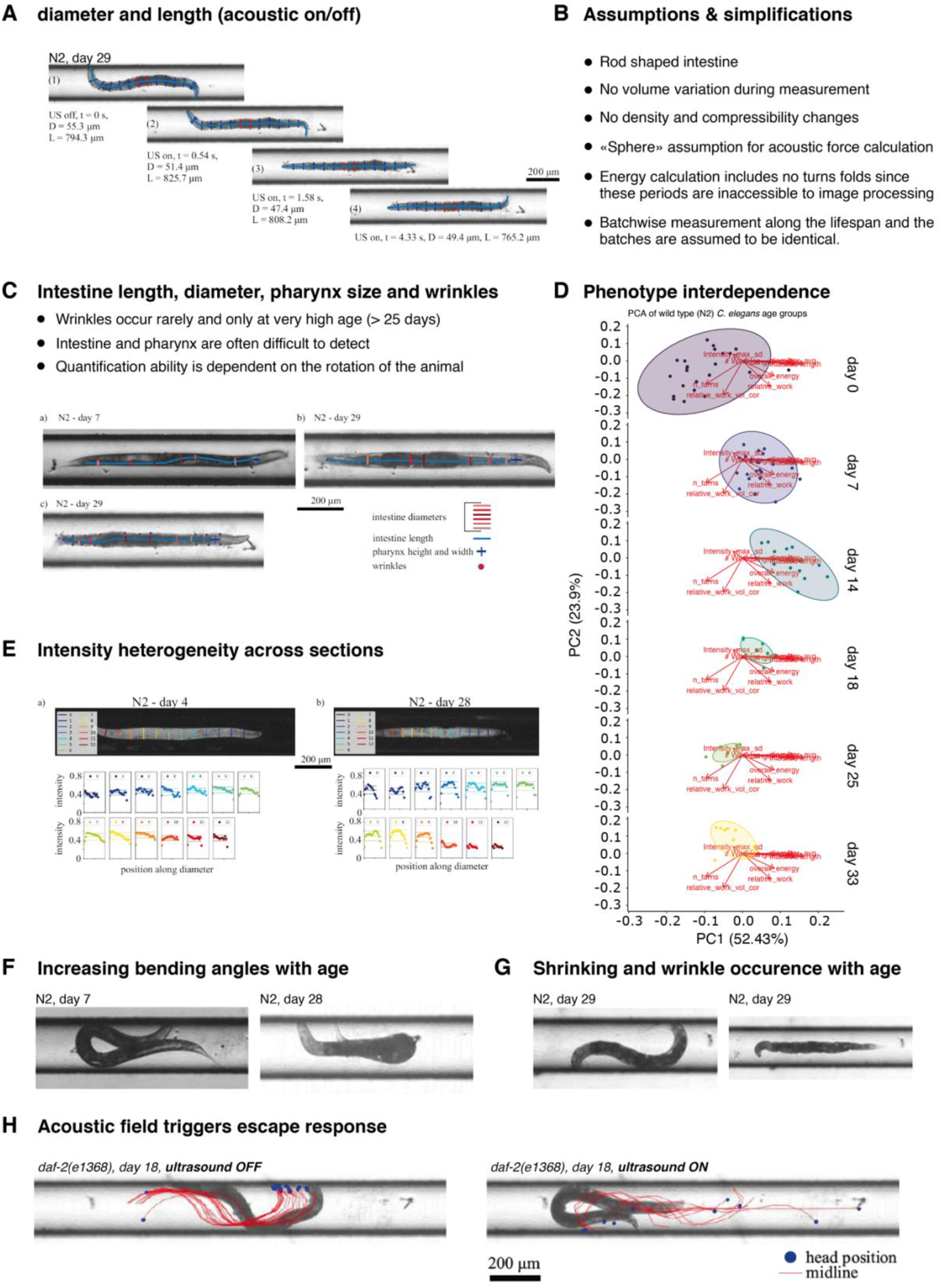
Phenotypes quantified within the microfluidic setup. Automated quantification of *C. elegans* diameter (D = diameter) evaluated at the middle segments colored in red and the length (L = length) of the animal along its spine colored in light blue (A), The change in diameter and length quantification as the field is switched (US = ultrasound) is indicated. The assumptions and necessary simplifications that were made to analyze the acquired frames are listed (B). Excerpts of the manual quantification of C. elegans intestine length and diameter, pharynx width and height, and the number of cuticle wrinkles are displayed (C). Annotation was performed in a self-developed image annotator suite based on the python programming language. Principal component analysis (PCA) of the measured phenotypes are displayed for aging wild type (N2) *C. elegans* individuals, with each point representing one individual. The overlaying components refer to the highly correlated features length, volume, and diameter. Tissue heterogeneity and age pigments were studied by analyzing the intensity distribution across different cross-sections along the animal’s fitted spine. Representative images of tissue and cuticle weakening displaying an increased bending angle of older animals compared to younger individuals as they attempt to turn in the channel (F). Similarly, older animals are smaller and sometimes display cuticle wrinkles (G). The strong stimulation of the acoustic force in the animals is shown for a representative *daf-2(e1368)* animal (H). The spline midline and head position is displayed for seconds preceding (left), and after (right), the acoustic field is activated. The animal directly responds with a strong escape response.

**Supplementary Figure 6.**
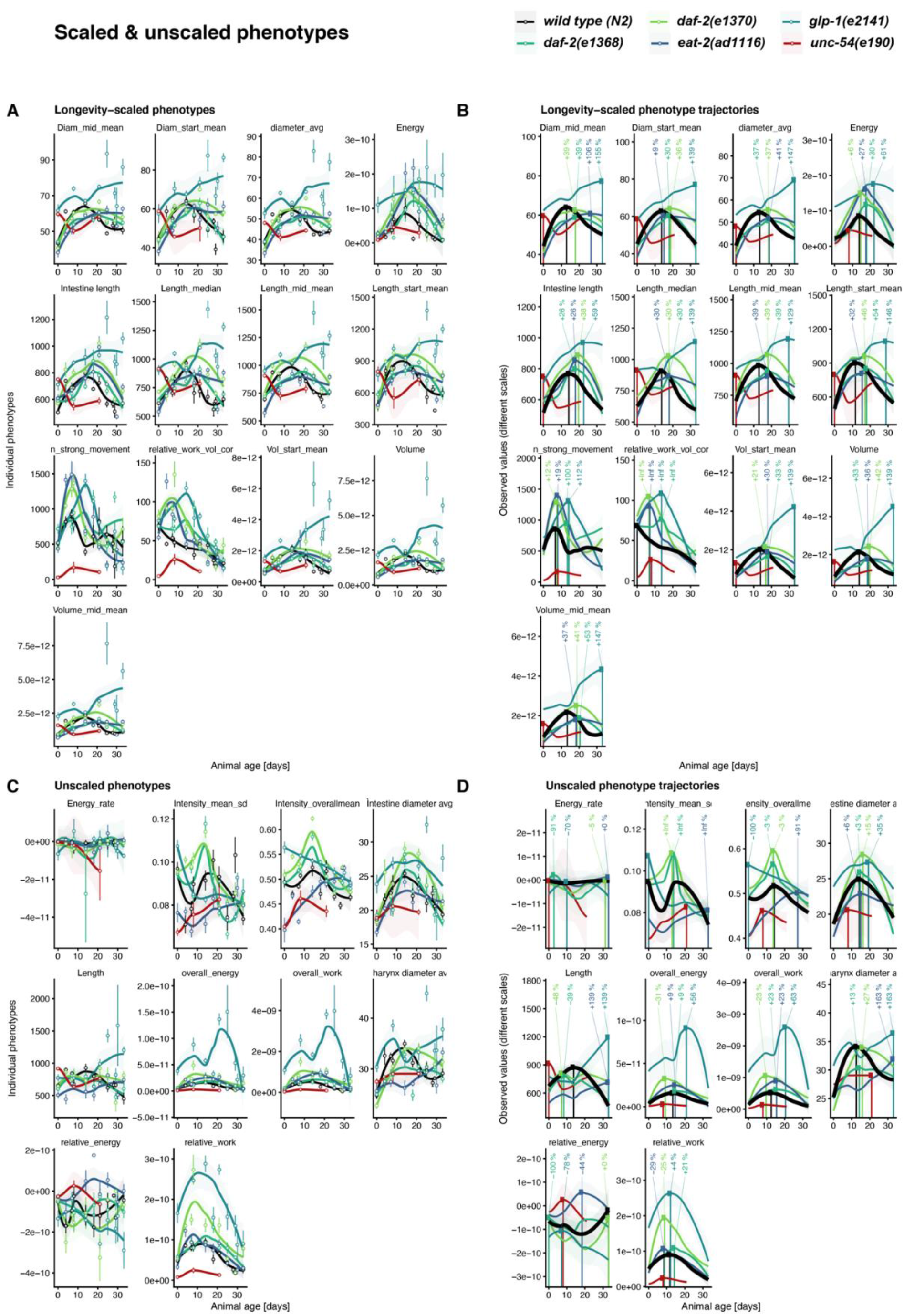
Rationale behind the partitioning of physical properties into scaled and unscaled phenotypes. Phenotypes displaying an age-dependent change and at least a partial temporal shift for any long-lived strain were manually classified as temporally scaled phenotypes (A, B), and phenotypes that do not satisfy these conditions are categorized as unscaled phenotypes (C, D). The quantified phenotypes are fitted using a loess model in all panels. The observed values for each individual are displayed using mean +/- standard error for each timepoint (A, C). The maximum value predicted by the loess fit is shown, and the increase relative to the wild type maximum is displayed at the top of the graph.

**Supplementary Figure 7.**
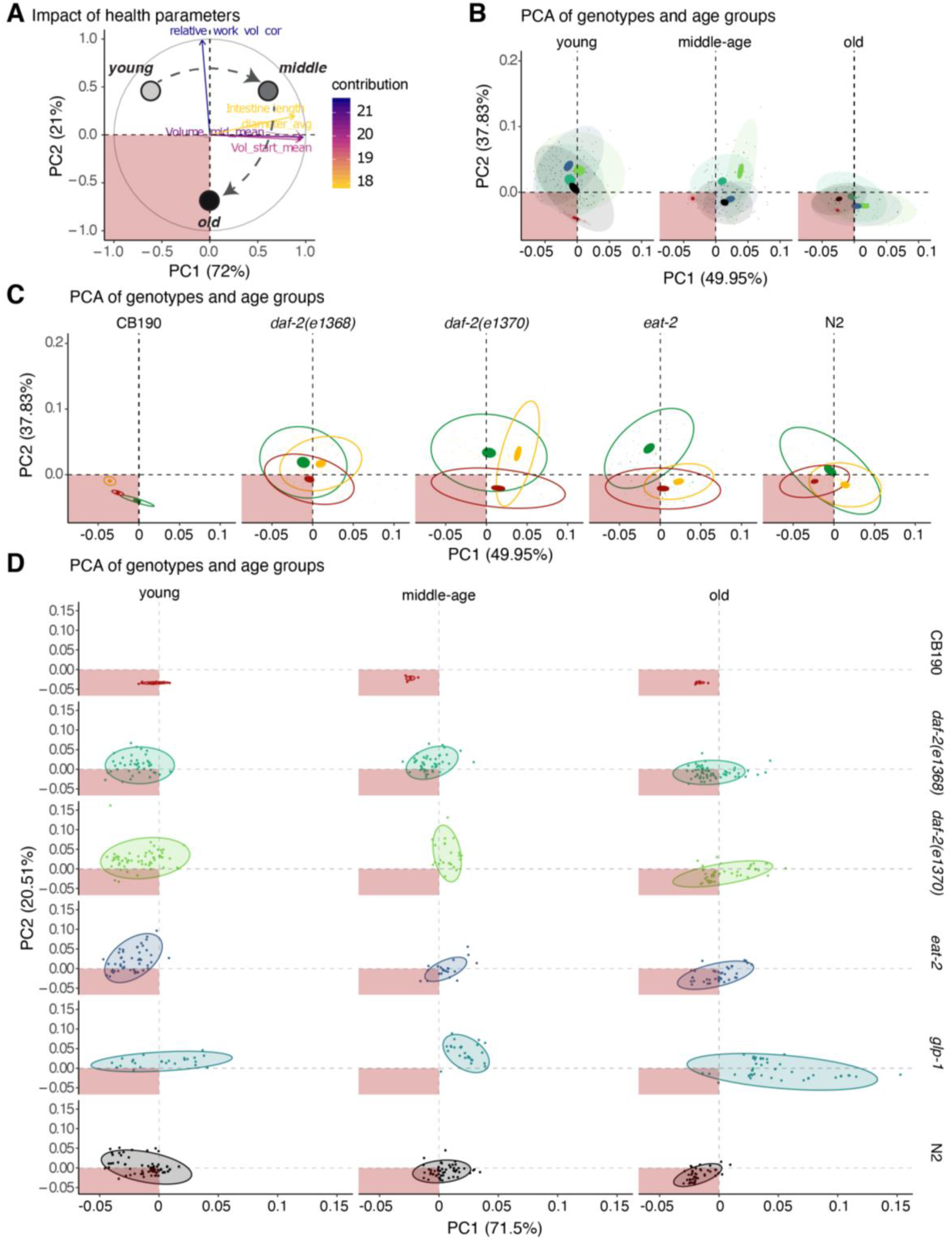
Principal component analysis of temporally scaled phenotypes to identify similarity between phenotypes and their contributions. The subset of temporally scaled phenotypes for each individual was subjected to principal component analysis, and the first two principle components are shown together with the variance they explain. The contribution of each phenotype to the two first principle components is displayed as vectors with the orientation reflecting the directionality and the color and length capturing the contribution of each phenotype (A). The overall progression of young to middle-aged to old individuals through the phenotype landscape is schematically illustrated. The bottom left quadrant is highlighted in red to indicate its association with animals experience a poor health status. To compare the effect of age on the clustering of the different *C. elegans* genotypes, all animals were grouped into 3 age categories, and the individual genotypes were shown in different colors (B). Confidence ellipses are drawn at the level of 95% for all samples (transparent) and the sample means (non-transparent). To compare the effect of age separately for each genotype, the three age categories are shown for each strain, young in green, middle-aged in yellow and old animals in red (C). Complete separation of age and genotype is provided in panel (D), highlighting the measurements for every individual animal encompassed by the 95% sample confidence ellipse.

**Supplementary Figure 8.**
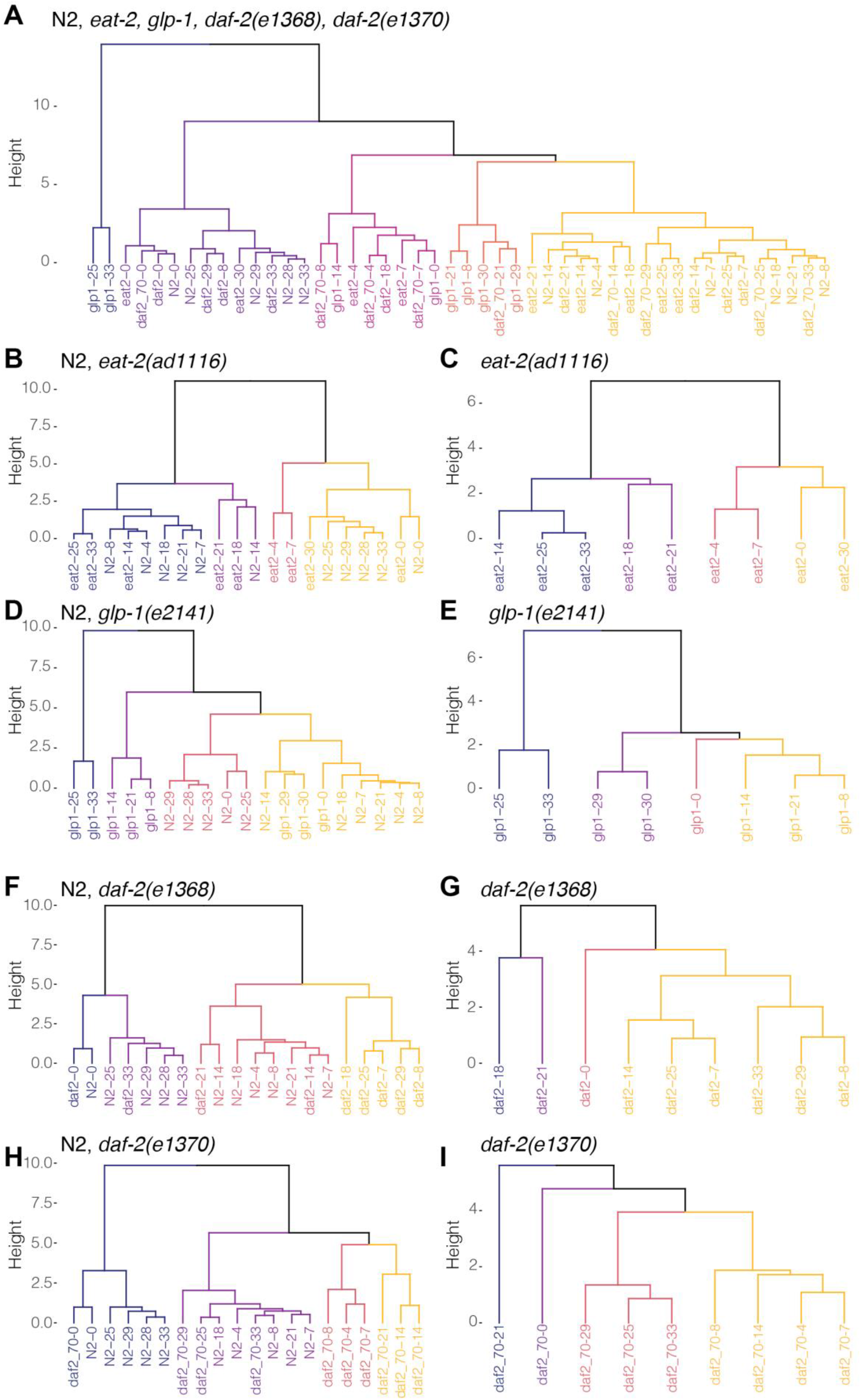
Hierarchical clustering of temporally scaled phenotypes. The similarity between sampled genotypes and ages are displayed using hierarchical clustering. The comparison between all samples (A), between each long-lived genotype and wild type (B, D, F, H), and within each long-lived is shown (C, E, G, I). The tree is cut into five clusters when comparing all samples and, in all other cases, four clusters and colored from left to right.

## Materials and Methods Strains

*Caenorhabditis elegans* strains were maintained on NGM plates and OP50 *Escherichia coli* bacteria. The wild-type strain was N2 Bristol. Mutant strains used are described at www.wormbase.org: LGII: *eat-2(ad1116)*; LGIII: *daf-2(e1368, e1370), glp-1(e2141)*.

### *C. elegans* culturing conditions

*C. elegans* populations were age-synchronized by isolating eggs from gravid C. elegans adults, incubating them for 18 hours in M9 until hatching in the presence of 5 μg mL-1 cholesterol (Sigma-Aldrich). Hatched L1-larvae were grown on standard culturing NGM OP50 plates and then shifted to 50 μM FUdR plates seeded with heat-killed OP50 plates when reaching the L4 state. Animals were maintained on FUdR plates until the measurement of different ages was taken. The *glp-1(e2141)* mutants were placed at 25°C after bleach treatment and shifted back to 15°C at the L4 stage and otherwise treated equally to the other strains. We note that there might be a possible additional lifespan extension from shifting *glp-1* larvae at 25°C during development since wild type N2 grown at 25°C and shifted to 20°C as adults showed increased lifespan (Zhang et al., 2015). Unfortunately, an overall same temperature regime for wild type and all mutants for this study is not possible since *daf-2* mutants would enter into dauer at 25°C during development. Thus, except for *glp-1* during its development, all animals were always maintained and aged at 15°C.

### Automated survival assays using the lifespan machine

To compare the lifespans among wild type and long-lived mutants, we raised all animals for several generations in parallel. Automated survival analysis was conducted using the lifespan machine setup described by (Stroustrup et al., 2013). Briefly, approximately 1000 L4 animals were resuspended in M9 and transferred to NGM plates containing 50 µM 5-Fluoro-2’deoxyuridine (FUdR) seeded with heat-killed OP50 bacteria and incubated at 15°C until day 4 of adulthood. Animals were then resuspended in M9 and transferred to fresh FUdR plates containing tight-fitting lids (BD Falcon Petri Dishes, 50×9mm), and the plates were dried with their lids open for 30 minutes after transfer. The plates were incubated for five additional days to rule out contamination and then loaded in the lifespan machine. Air-cooled Epson V800 scanners were utilized for all experiments operating at a scanning frequency of one scan of 30 minutes. Temperature probes (Thermoworks, Utah, US) were used to monitor the temperature on the scanner flatbed and kept constant at 15°C.

For the health-, sick-, life-span validation experiment, we chose to alter three experimental conditions: 1. temperature regime, 2. different bacterial source and live bacteria, and 3. no FUdR. We maintained temperature-sensitive sterile mutants TJ1060 *spe-9(hc88)* I; *rrf-3(b26)* II and *glp-1(e2141)* at 15°C. Synchronized L1 by bleach preparation and let the larvae develop to day-2 adults on OP50 NGM plates at 25°C. Then, we transferred worms onto empty vector L4440 HT115 bacteria at 20°C and at day-8 of adulthood to lifespan plates containing L4440 bacteria to assess lifespan at 20°C.

### Voluntary movement healthspan measured by lifespan machine

The time point at which the animal stops moving completely and irretrievably is classified as the point of death and defines the lifespan of each individual. The health-to sickspan transition is estimated by the time point when major movement ceases, and exclusively head movements, posture change, and minor body movements can be observed. The animal is also required to be sedentary and remains in the rough vicinity of the area; it will ultimately die.

### Microfluidics device measuring muscle strength

A detailed description of the microfluidics device development and characterization is found in the *Cell Reports Methods* manuscript.

## Figure generation and statistics

The analysis was performed using the statistical software R. data processing and visualization were performed using the tidyverse package collection, most prominently dplyr and ggplot2. Furthermore, packages were used for lifespan analysis (survival, survminer), computing and visualizing PCA (stats and ggfortify, factoextra), fitting loess models (stats), and segmented fits (segmented), labeling (ggrepel) comparing distributions (ggpubr), and arranging figures (cowplot).

